# CZ ID: a cloud-based, no-code platform enabling advanced long read metagenomic analysis

**DOI:** 10.1101/2024.02.29.579666

**Authors:** Sara E. Simmonds, Lynn Ly, John Beaulaurier, Ryan Lim, Todd Morse, Sri Gowtham Thakku, Karyna Rosario, Juan Caballero Perez, Andreas Puschnik, Lusajo Mwakibete, Scott Hickey, Cristina M. Tato, CZ ID Team, Katrina Kalantar

**Affiliations:** Chan Zuckerberg Initiative, Redwood City, California, USA; Oxford Nanopore Technologies Inc., New York, NY, USA; Chan Zuckerberg Biohub San Francisco, San Francisco, California, USA; The Bioinformatics CRO, Orlando, FL, USA

**Keywords:** Nanopore, mNGS, microbiome, divergent virus, novel virus, non-human host

## Abstract

Metagenomics has enabled the rapid, unbiased detection of microbes across diverse sample types, leading to exciting discoveries in infectious disease, microbiome, and viral research. However, the analysis of metagenomic data is often complex and computationally resource-intensive. CZ ID is a free, cloud-based genomic analysis platform that enables researchers to detect microbes using metagenomic data, identify antimicrobial resistance genes, and generate viral consensus genomes. With CZ ID, researchers can upload raw sequencing data, find matches in NCBI databases, get per-sample taxon metrics, and perform a variety of analyses and data visualizations. The intuitive interface and interactive visualizations make exploring and interpreting results simple. Here, we describe the expansion of CZ ID with a new long read mNGS pipeline that accepts Oxford Nanopore generated data (czid.org). We report benchmarking of a standard mock microbial community dataset against Kraken2, a widely used tool for metagenomic analysis. We evaluated the ability of this new pipeline to detect divergent viruses using simulated datasets. We also assessed the detection limit of a spiked-in virus to a cell line as a proxy for clinical samples. Lastly, we detected known and novel viruses in previously characterized disease vector (mosquitoes) samples.

## INTRODUCTION

Metagenomic next-generation sequencing (mNGS) is a powerful approach that uses sequencing technologies to comprehensively analyze a sample’s genetic material, including host-associated microbes (e.g., bacteria, viruses, fungi, and parasites).

Metagenomics has emerged as an effective lens for studying infectious diseases, enabling the unbiased, direct detection and identification of pathogens from clinical (e.g., Chiu and Miller, 2019; Li *et al*., 2021; Bohl *et al*., 2022), non-human host (e.g., Batson *et al*., 2021; Ergunay *et al*., 2022; Juergens *et al*., 2022), and environmental samples (e.g., Datta *et al*., 2020; Farrell *et al*., 2022; Ramuta *et al*., 2023; Urban *et al*., 2023). Unlike traditional culture-independent methods that rely on targeted assays to identify microbes, metagenomics can potentially detect all microbes present in a sample, regardless of whether they have been previously characterized. This feature is vital for identifying new or emerging pathogens that may be missed using traditional methods. Metagenomics can also detect novel variants of known pathogens, which is essential for tracking the spread of drug-resistant pathogens or identifying emerging strains. Metagenomics can provide quick results where unsuspected pathogens threaten public health by providing a complete view of microbial composition. For example, in the case of an outbreak or epidemic, metagenomics can rapidly identify and characterize the causative agent to inform public health interventions and help prevent further spread of the disease (e.g., Wu *et al*., 2020).

Short read sequencing technologies are widely used in metagenomics, but long read sequences have two potential advantages for mNGS. First, long reads allow for more accurate and comprehensive *de novo* assembly of metagenomes, providing a detailed view of the microbial community in a given sample (Portik *et al*., 2022). Second, long reads enable the characterization of genomic structural variants (i.e., insertions, deletions, and rearrangements) that are difficult to identify using short reads (Mahmoud *et al*., 2019). Since structural variants can be determinants of pathogen virulence and antibiotic resistance, their accurate identification has implications for clinical decision-making (e.g., Dai *et al*., 2022; Schikora-Tamarit and Gabaldón, 2022). Long read sequencers like the portable MinION from Oxford Nanopore Technologies are also comparatively small, affordable, and easy to maintain (Yek *et al*., 2022), lowering the barrier to obtaining high-quality long reads and democratizing these benefits.

However, considerable technical and computational challenges are associated with processing and analyzing large, complex datasets generated by long read sequencing platforms. Frequently, processing mNGS datasets requires expert bioinformatic skills, access to powerful computers, and long runtimes. These issues hinder the use of metagenomics in infectious disease research, especially in resource-limited settings (Yek *et al*., 2022; Marais *et al*., 2023). Here, we introduce a new metagenomics module of the CZ ID platform that analyzes long read data from Oxford Nanopore Technologies to address data analysis challenges. The CZ ID mNGS Nanopore module provides infectious disease researchers with a fast, accurate, and free tool for processing long read data and characterizing complex microbial communities without the need for coding or computing resources. Using microbial community standards, simulated datasets, and real-world samples, we demonstrate the potential of the CZ ID mNGS Nanopore pipeline. We highlight the pipeline’s ability to identify known and novel viruses from clinical and non-human host samples.

## IMPLEMENTATION

### CZ ID mNGS Nanopore pipeline

CZ ID is a free, cloud-based genomic analysis platform that enables researchers to detect pathogens using metagenomic data, predict antimicrobial resistance genes, and generate viral consensus genomes (Kalantar *et al*., 2020). We describe the Nanopore metagenomics module (v0.7). For up-to-date information, please see the documentation at https://chanzuckerberg.zendesk.com/hc/en-us. All of the code is open-source and available at https://github.com/chanzuckerberg/czid-workflows/tree/main/workflows/longread-mngs.

The CZ ID mNGS Nanopore pipeline accepts basecalled Oxford Nanopore reads from DNA or RNA samples sequenced from any host organism or environment in FASTQ format (compressed and uncompressed). Reads and associated metadata can be uploaded via the CZ ID web application (https://czid.org/) or command line interface (https://github.com/chanzuckerberg/czid-cli/). To account for unexpected error rates, users need to specify which ONT basecalling model was used to generate the data (i.e., fast, hac, sup).

Users upload FASTQ files to CZ ID directly from their computers. Sequence files are automatically concatenated during upload if there are multiple FASTQ files associated with the same sample. The platform recognizes these files by the base name including the qualifiers “_pass_” or “fastq_runid_”. To add metadata, users can enter information directly through the web interface or upload a metadata file in CSV format. There are six required metadata fields: Host Organism, Sample Type, Water Control, Nucleotide Type, Collection Date, and Collection Location.

Once data is uploaded, the CZ ID mNGS Nanopore pipeline is executed in the cloud on Amazon Web Services (AWS) infrastructure. The pipeline workflow consists of four major steps:1) host and quality filters, 2) *de novo* assembly, 3) alignment to NCBI, and 4) taxon reporting (Fig. 1).

**Figure 1.**
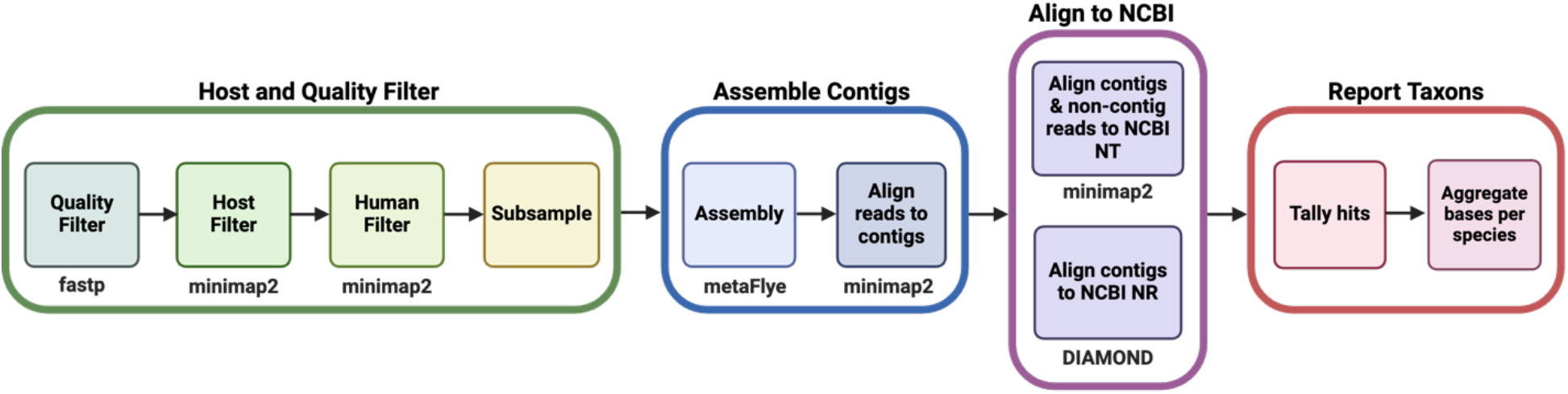
Overview of the CZ ID Nanopore mNGS pipeline.

### QC and host filtering

CZ ID executes quality control steps with the program fastp (Chen *et al*., 2018) to remove reads of low quality (mean phred score < 9), low complexity (< 30%), and short length (< 100 bp). CZ ID then filters out host DNA by aligning all reads to a host reference genome (specified during upload) using minimap2 (Li, 2018). Regardless of the host species, all reads that map to *Homo sapiens* are removed to eliminate possible human contamination that may have occurred during sample preparation. Samples with a high fraction of non-host reads (e.g., stool samples) could retain large numbers of sequences following host- and quality-filtration steps. Therefore, non-host reads are subsampled to 1 million reads before proceeding to *de novo* assembly to control the computational time and cost.

### Assembly based alignment

Non-host reads then undergo *de novo* assembly for two reasons: 1) to improve precision and sensitivity during mapping to reference databases and 2) to enable the recovery of metagenome-assembled genomes (MAGs). Long reads (> 1000 bp) are assembled using metaFlye (Kolmogorov *et al*., 2020), an algorithm in the Flye assembler (v2.9.2) developed for long read metagenomes (Kolmogorov *et al*., 2019). If reads are basecalled with the Guppy super accuracy model (sup), then the Flye option “--nano-hq” is applied; otherwise, “--nano-raw” is used and paired with one round of polishing (within Flye) following assembly. These flags are intended to enable optimization for different error rate profiles. During the assembly step, the metaFlye output loses the information about which reads belong to each contig. Therefore, CZ ID uses minimap2 to map the subsampled non-host reads to the assembled contigs and SAMtools (Li *et al*., 2009) to extract non-contig reads. Read mapping information is then used to count the number of reads and bases that map to each contig and calculate coverage statistics for all assembled contigs.

### Taxon reporting

To assign a taxonomic identity to each contig, CZ ID maps contigs to the NCBI nucleotide (NT) and non-redundant protein (NR) databases using minimap2 and DIAMOND (Buchfink *et al*., 2015), respectively. Non-contig reads are identified by mapping to the NT database. However, higher error rates in non-contig Nanopore reads preclude their use for NR alignment, given the need to translate to amino acid sequences accurately. Any reads that fail to map to the databases are removed, compiled into an unmapped_reads file, and made available to download for further analysis. As an additional host-filtering measure, for any host that is a Deuterostome, hits to the NT database matching GenBank accessions in the superphylum *Deuterostomia* are removed, given the high likelihood that such reads are of host origin. All hits that match artificial sequences (NCBI:txid81077) are also removed from the sample report. Contigs aligning to NT and NR NCBI accessions are assigned the corresponding taxonomic identifiers (taxIDs). If contigs align equally well to multiple taxa, then a single taxID is randomly selected. Reads assembled into contigs are assigned the same taxID as their parent contig. Finally, results are aggregated to produce NT and NR counts for each taxID at both the species and genus levels. Each pipeline run is versioned, showing the database index version used in the platform.

The CZ ID web app provides individual sample results that can be explored in the Sample Report (Fig. 2), an interactive table summarizing identified taxa and match metrics, including bases per million (bPM), bases (b), reads (r), contig, contig bases (contig b), percent identity (%id), length (L), and expect value (E value) (Table 1).

**Table 1.** Metrics and definitions are reported per taxa in the CZ ID web app Sample Report. *All metrics are reported for alignments against the NCBI nucleotide (NT) and non-redundant protein (NR) databases separately. Note that values against NR only reflect alignments between contig sequences and their matching taxon (i.e., unassembled reads are not aligned against NR).

| Metric* | Definition |
| --- | --- |
| bPM | **Bases per million** - Number of bases within all the reads aligning to a given taxon, including those assembled into contigs that mapped to the taxon, per million bases sequenced. |
| b | **Bases** - Number of bases within all the reads aligning to a given taxon, including those assembled into contigs that mapped to the taxon. |
| r | **Reads** - Number of reads aligning to a given taxon, including those assembled into contigs that mapped to the taxon. |
| contig | **Contigs** - Number of assembled contigs aligning to a given taxon. |
| contig b | **Contig bases** - Number of bases within all the reads that assembled into contigs aligning to a given taxon. |
| %id | **Percent identity** - Average percent identity between all the query sequences (contigs and unassembled reads) and their matching taxon. |
| L | **Length** - Average length of alignments between all the query sequences (contigs and unassembled reads) and their matching taxon. Note that values against NR are reported in base pairs. |
| E value | **Expect value** - Average expect value (E-value) of alignments against the NT and NR databases. The E-value represents the number of matches with similar quality one would “expect” to see by random chance. This parameter provides a measure of randomness. The lower the E-value, the lower the probability of getting a match or alignment by random chance (i.e., E-value tends towards 0 for significant alignments). |

**Figure 2.**
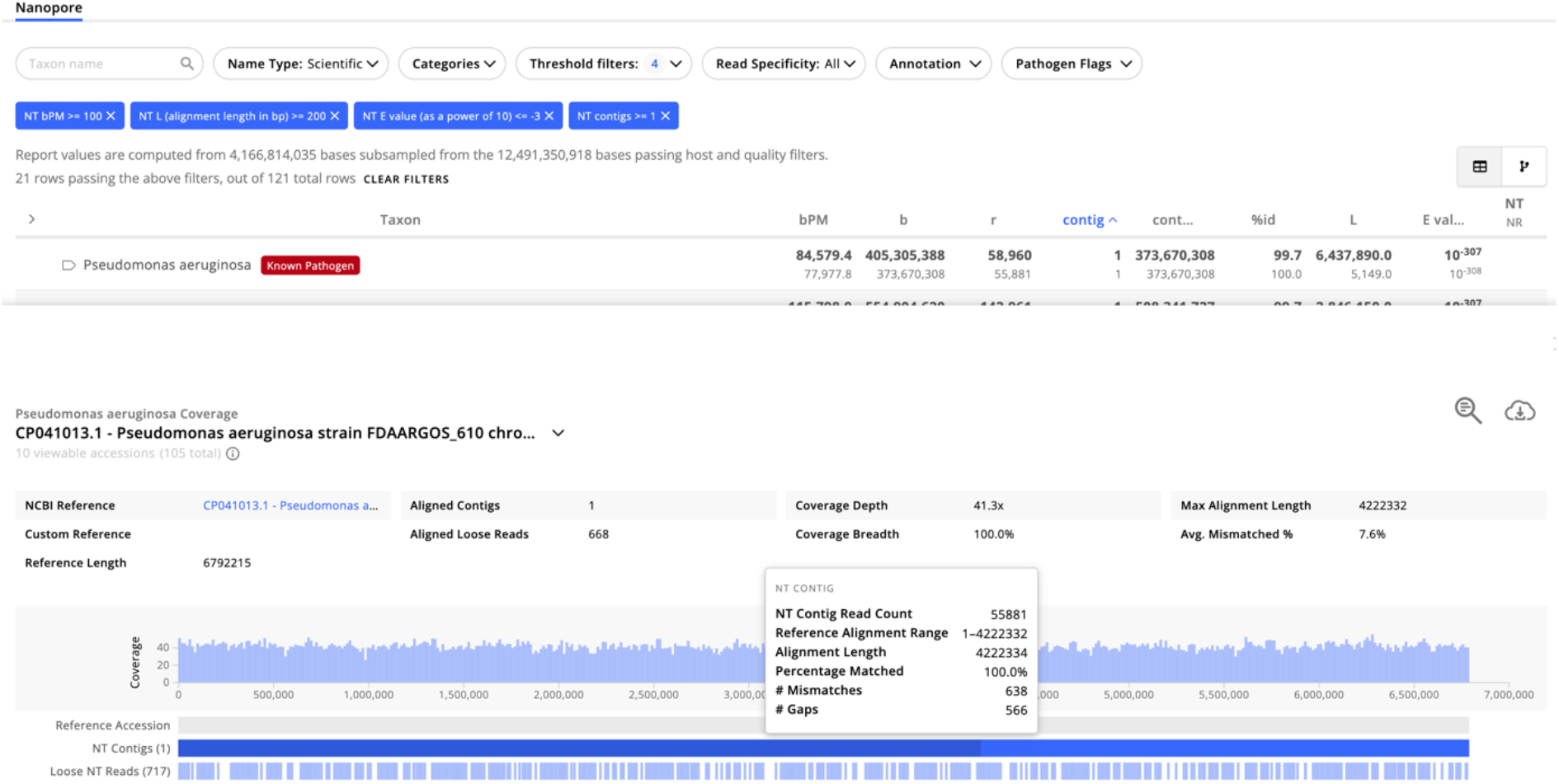
CZ ID web app Sample Report with a coverage visualization for a *Pseudomonas aeruginosa* hit showing the coverage plot and statistics including coverage depth, breadth, max alignment length and average percent mismatched. The rows below the coverage plot with blue bars represent contig and non-contig (loose) reads aligned to the reference accession on the NCBI NT database.

### Sample report filters and thresholds

Since metagenomic analysis is a non-targeted approach that captures microbial composition, the number of taxa on the CZ ID Sample Report can be very large, and not all reported taxa may be relevant to every research question. Filtering the results can help focus on abundant species representing microbial groups of interest. To do this, users can filter the Sample Report by category based on specific microbial groups, including Archaea, Bacteria, Eukaryota, Viroids, Viruses (all viruses), Viruses - Phage (only phage), and Uncategorized (not assigned to a specific taxonomic group).

Organisms with known human pathogenicity are tagged in red. The list of organisms with known pathogenicity recognized in CZ ID is available here: https://czid.org/pathogen_list.

In addition to category filters, users can set threshold filters to remove spurious matches based on metric value ranges reflecting the quality of alignments against NT or NR databases (Table 2). For example, to identify high-confidence hits to microbes, set a bPM > 100 filter for matches in the NT database to remove taxa that were present at low levels; set a bPM > 1 filter for matches in the NR database to remove taxa that only have matches in non-coding regions (i.e., taxa that only have matches in the NT database); set a L > 200 bp filter for matches in the NT database to remove taxa for which alignments were < 200 bp. The longer the alignment between a query sequence and its matching taxon, the greater confidence users can have in the match. Set an E-value < 0.001 filter for matches against NT and NR to remove likely random matches. Since the E-values are specified as a power of 10, specify an E-value ≤ -3 when setting this filter. The lower the E-value, the more stringent the filter will be. We suggest setting the E-value filter to ≤ -10 for stringent searches or ≤ -1 to allow for less significant taxon matches. These thresholds are suggestions based on typical sources of error in metagenomic analysis but can be adjusted based on the needs of studies and sample types.

**Table 2.** Suggested threshold settings to filter out spurious matches to NCBI NT or NR databases.

| Criteria | Threshold filter | Value |
| --- | --- | --- |
| Filter for abundant taxa | NT bPM | $\geq 100$ |
| Remove hits to only non-coding regions | NR bPM | $\geq 1$ |
| Remove short alignments (potentially low-confidence due to shared homology) | NT L | $> 200$ bp |
| Remove random matches | NT/NR E-value | $< -3$ |

Divergent or novel sequences may have matches at the amino acid level but not at the nucleotide level. If a taxon has high NR counts and low or no NT counts, it is likely a novel sequence or an organism that does not have sequences in the NCBI databases. Novel sequences are more likely to match the protein database since amino acid sequences are more conserved across taxa than nucleotide sequences. In our experience, this pattern is more commonly observed for viral sequences compared to bacterial sequences.

### Visualizations

There are three types of visualizations associated with CZ ID’s mNGS Nanopore module, including the taxonomic tree view, coverage, and pipeline visualizations. Users can explore an overview of all detected species using Sample Report (Fig. 2) and the Taxonomic Tree View (Fig. 3). The tree view depicts the taxonomic lineages of all microbes identified in a given sample. The weight or thickness of the lines connecting tree nodes is proportional to the metric selected to visualize the tree. By default, the tree lines will reflect values representing the number of bases (b) matching a given taxon in the NT database (NT b). Taxa with thicker lines will have proportionally higher NT b values than those with lower values.

**Figure 3.**
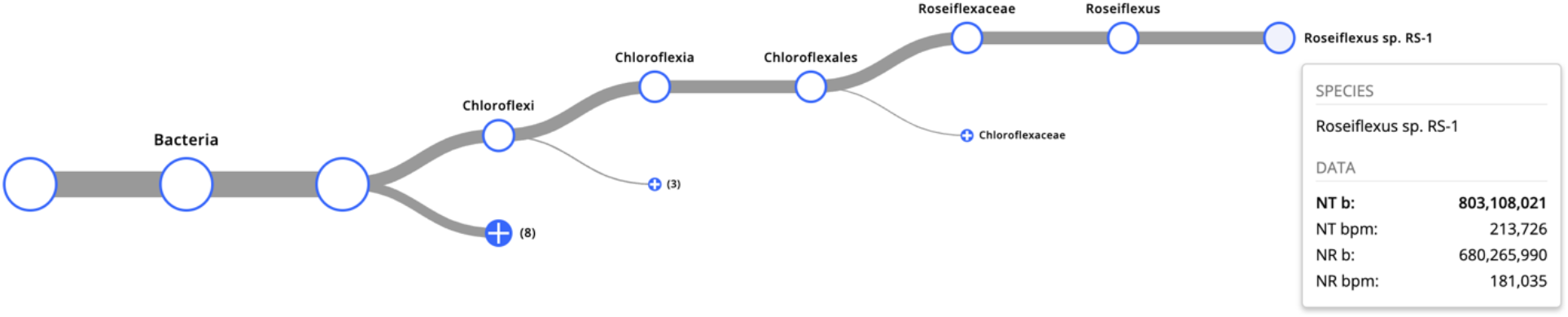
Taxonomic tree view of the CZ ID web app Sample Report showing taxonomic hits in a cladogram.

The sample report includes a coverage visualization to examine the uniformity and breadth of genome coverage for a taxon of interest (Fig. 2). This feature is available for all taxa supported by at least one read matching at the nucleotide level (NT database).

CZ ID also provides a detailed interactive visualization of the pipeline steps implemented for each sample, enabling users to find details about each step and download intermediate files of interest. For example, by navigating the pipeline visualization to select the step for “Unmapped Reads”, users can download reads that did not align to NCBI NT and NR databases (i.e., unmapped reads).

## METHODS AND MATERIALS

### Benchmarking: dataset, tools, and metrics

To benchmark CZ ID’s performance in detecting known microbes against other tools, we used a previously published microbial community standard (Nicholls *et al*., 2019), ZymoBIOMICS Microbial Community Standard (Zymo Research Irvine, CA, USA, Product D6300, Lot ZRC190633), basecalled using Guppy v6.0.1 with the super accuracy model. ZymoBIOMICS Microbial Community Standard is a commercially available reference standard comprising ten microbial species at known relative abundances.

We uploaded rebasecalled reads to CZ ID, chose human as the host, ran the pipeline, and downloaded the sample report. Next, we compared CZ ID NT results against Kraken2 v2.1.3 (Wood *et al*., 2019), a widely used open-source bioinformatic tool for analyzing metagenomic data. Kraken2 results were generated by aligning the filtered and subsampled reads (1 million) from CZ ID against the Kraken2 Standard plus RefSeq protozoa & fungi database (downloaded 10/9/2023). Finally, we computed precision and recall, generated the precision-recall curves, and estimated the AUPR and L2 distance, as detailed in (Ye *et al*., 2019).

### Application I: Detecting divergent viruses

One of the most compelling applications of mNGS is the identification of divergent or novel pathogens, as illustrated by the fact that SARS-CoV2 was first sequenced using metagenomics (Wu *et al*., 2020). Viruses have high mutation rates and can evolve over short timescales (Peck and Lauring, 2018). Therefore, known viruses may diverge from reference sequences currently in databases. Identifying and tracking viruses as new variants evolve is essential for virologists, epidemiologists, and public health officials (Grubaugh *et al*., 2019). Here, we evaluated the sensitivity of the CZ ID mNGS Nanopore pipeline in detecting known variants across diverse virus families and varying genome sizes (Suppl. Table 1). We performed *in silico* evolution of six virus species from the reference sequence from 5% to 50% divergent at the nucleotide level (scripts are available at https://github.com/caballero/mutator/). We simulated Nanopore reads from each at 7X depth using PBSIM2 (Ono *et al*., 2021), ran them through the CZ ID platform using parameters for “hac” Guppy basecaller setting and “ERCC only” host, and evaluated the divergence thresholds at which CZ ID could detect the expected viral species.

### Applications II and III: Identification of microbes in human clinical and complex non-human host samples

It is crucial to detect known, divergent, and novel viruses in clinical and complex environmental samples, especially RNA viruses. Many of the viruses that currently cause severe disease in humans (e.g., COVID-19, dengue, influenza, measles, polio, AIDS, chikungunya, Ebola, rabies, and Lassa fever) have RNA genomes. RNA viruses can evolve new variants quickly, mutating away from sequences in reference databases. RNA viruses are the primary infectious pathogens of emerging (∼44%) and novel (∼66%) human infectious diseases (see Cassarco-Hernandez *et al*., 2017 for a review). Moreover, the vast majority of human-infecting RNA viruses are considered zoonotic in origin (89%), highlighting the importance of their identification in a broad range of host organisms (Woolhouse *et al*., 2013).

To simulate a human clinical sample and evaluate the sensitivity of CZ ID for detection of a known spiked-in virus, HeLa cells were infected with human coronavirus OC43 (ATCC, #VR-1558) at varying multiplicity of infection (MOI) values (MOI = 1, 0.1, 0.01, 0.001, 0.0001, no virus), and incubated for 24 hours prior to collection and storage in RNA shield. Nucleic acid was extracted using the quick-DNA/RNA Pathogen MagBead kit (Zymo Research). Extracted nucleic acid was treated with DNAse to isolate RNA and run on a TapeStation (Agilent) for quality control to examine RNA integrity.

To test whether CZ ID can characterize the virome of non-human hosts (e.g., disease vectors), we selected five mosquitos that had been previously collected, screened using Illumina, and analyzed in CZ ID (Batson *et al*., 2021). Details on how the individual mosquitoes were collected and stored and how RNA was extracted can be found in the methods described by Batson *et al*., (2021).

We then used 10 ng of extracted RNA from each sample (HCoV OC43 and six mosquito samples; Supp. Table 2) as input to a variation on the SISPA (Sequence-Independent Single Primer Amplification) protocol detailed below (Claro *et al*., 2021).

### Library preparation of RNA for cDNA sequencing

We prepared the RT primer mix by adding 19 μl of RLB RT 9xN primer (10 μM), 1 μl of RLB 15xT poly(dT) primer (10 μM), and 80 μl of nH_2_O to a 1.5 ml tube with a final concentration of 2 μM. Next, we adjusted RNA sample (10 ng) volumes accordingly with up to 10 μl of nH_2_O to each tube, followed by 1 μl of the RLB RT 9N/15xT primer mix (2 μM) and 1 μl of the dNTP mix (10 mM). Samples were incubated at 65 °C for 5 mins, followed by snap-cooling on a pre-cooled PCR block for 2 mins.

For each reaction, we mixed 12 μl of annealed RNA from the previous step, 4 μl of Maxima H(-) buffer (5X), 1 μl of RNase OUT, and 2 μl of TSOmG (2 μM) and incubated at 42 °C for 2 mins. Next, we added 1 μl of Maxima H(-) enzyme to each reaction, resulting in a final volume of 20 μl. We incubated the samples at 42 °C for 90 mins and subsequently at 80 °C for 5 mins to complete the reverse transcription and cDNA synthesis. To fragment the library, we added 1 μl of FRM to all tubes, incubated at 30 °C for 1 min and then at 80 °C for 1 min and cooled the samples on ice.

### PCR amplification

The total PCR reaction volume was 50 μl, with 5 μl of tagmented cDNA from the previous step, 25 μl of LongAmp Taq 2X master mix, 1 μl of RLB 01-12 (10 μM), and 19 μl of nH_2_O. We performed the PCR protocol with an initial denaturation step at 95 °C for 45 s, followed by denaturation at 95 °C for 15 s, annealing at 56 °C for 15 s, and extension at 65 °C for 6 mins. The final extension step was at 65 °C for 10 mins, and we held the reaction at 10 °C.

### Cleanup and quantification

To clean up the PCR products, we added 1 μl of Exonuclease I to each reaction and incubated the samples at 37 °C for 15 mins, followed by 80 °C for 15 mins. We used AMPure XP beads for purification. After resuspending the beads by vortexing, we added 0.8x SPRI beads to the samples. The mixtures were then incubated for 5 mins on a rotator mixer to facilitate binding of DNA to the beads. After incubation, we pelleted the samples on a magnet and carefully removed the supernatant. We washed the beads twice with freshly prepared 80% ethanol in Nuclease-free water. For elution, we added 15 μl Elution Buffer (EB) to the pellet and then discarded the beads. We quantified the DNA library using a Qubit and determined the library size by running 1 μl on a 12k Agilent Bioanalyzer.

### Library preparation for barcoded DNA and Nanopore sequencing

To prepare the barcoded DNA for sequencing, we made up the sample to 50 fmols in 11 μl of EB. Next, we added 1 μl of RAP-F to the barcoded DNA. The tube was gently flicked to mix the contents, and then we spun it down. The reaction was incubated for 5 mins at room temperature. Libraries were then sequenced on a GridION using R9.4.1 flowcells, and raw data was basecalled with Guppy v5 (sup).

### Analysis (mosquito virome)

Raw reads were uploaded to CZ ID and analyzed using NCBI databases (NT, NR) dated prior to the publication of Batson *et al*., (2021). We downloaded the results from CZ ID and filtered for hits with ≥ 1 contigs matching to NT or NR databases. All hits were further investigated using BLASTn, either directly through CZ ID (NT contigs) or by downloading the contigs associated with the NR hit and using NCBI BLASTn to identify the top hit. We then compared the top hits for viruses to those found in Batson *et al*., (2021), noting whether each virus was labeled as “novel” or “known”.

## RESULTS

### Benchmark of microbial community standard

We analyzed the relative abundance of a mock microbial community standard to benchmark CZ ID’s performance in detecting known microbes (bacteria and fungi) and compared it against Kraken2, another commonly used metagenomics tool. For each tool, we computed the AUPR and L2 distance. The relative abundance estimates for CZ ID NT and Kraken2 were nearly identical (Fig. 4), with the values for both statistics being equivalent (AUPR = 1.0 and L2 distance = 0.7 for both tools).

**Figure 4.**
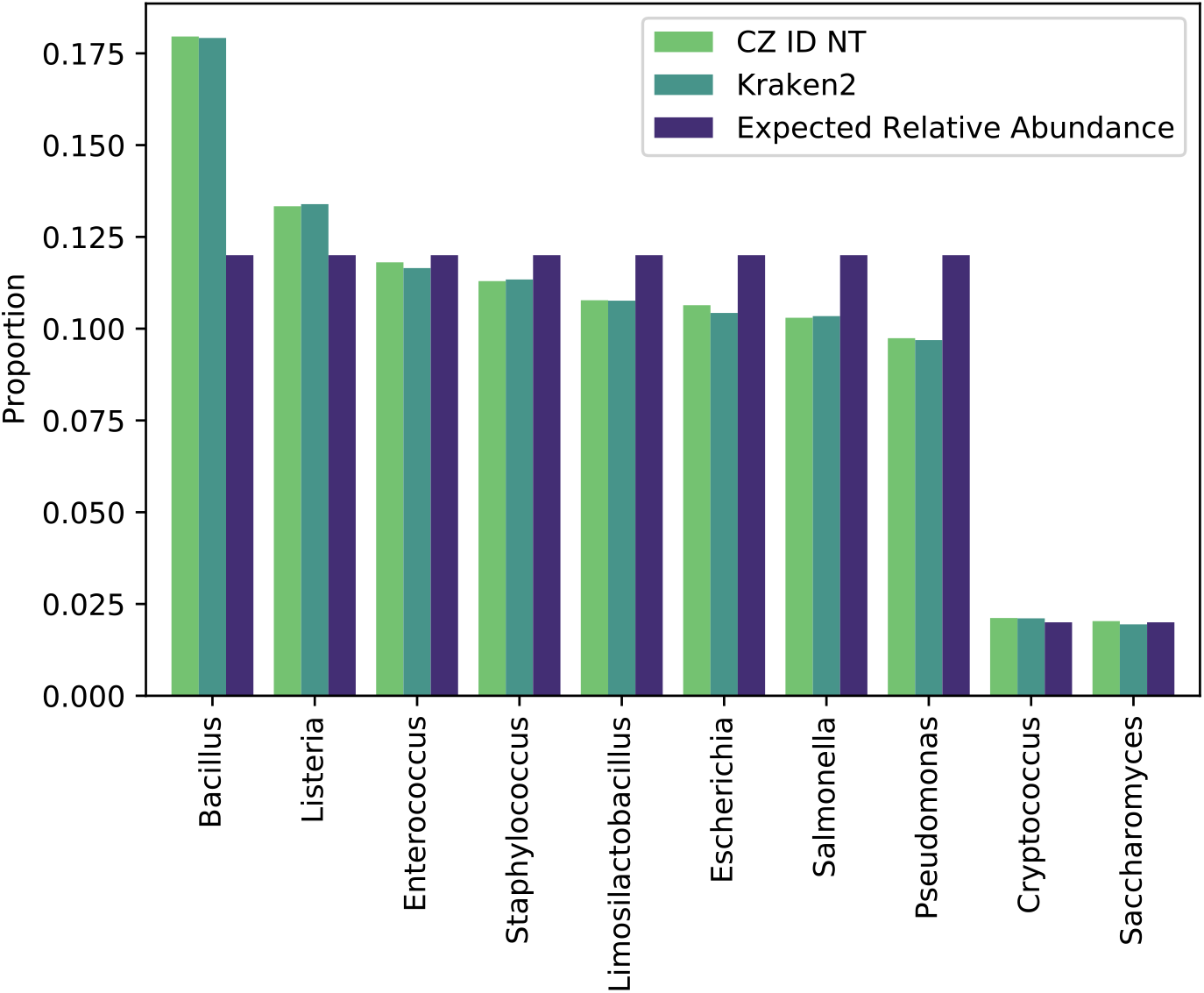
Relative abundance estimates (proportion) of 10 known microbial genera. The expected relative abundance values were based on the known abundance in the mock sample published by the manufacturer.

Beyond relative abundance estimates, the CZ ID pipeline produces assemblies that can provide additional value. In this sample, CZ ID *de novo* assembled metagenome-assembled genomes (MAGs) for four of the eight bacterial species in the mock community. Specifically, CZ ID assembled a complete MAG in a single contig for *Pseudomonas aeruginosa* (6,791,196 bp; 41X) and two contigs for *Listeria monocytogenes* (3,006,941 bp; 136X) and *Limosilactobacillus fermentum* (1,904,687 bp; 162X). For *Escherichia coli*, CZ ID assembled a complete MAG (2 contigs; 4,764,698 bp; 65X) and a plasmid (1 contig; 56X).

### Application I: Detecting divergent viruses

We evaluated the sensitivity of CZ ID for detecting divergent variants of six known viruses. The results for NT showed that CZ ID detected viruses up to 10-20% nucleotide sequence divergence from the reference genomes (or 80-90% similar) (Fig. 5). Results for NR had higher sensitivity, detecting viruses with 40-50% sequence divergence.

**Figure 5.**
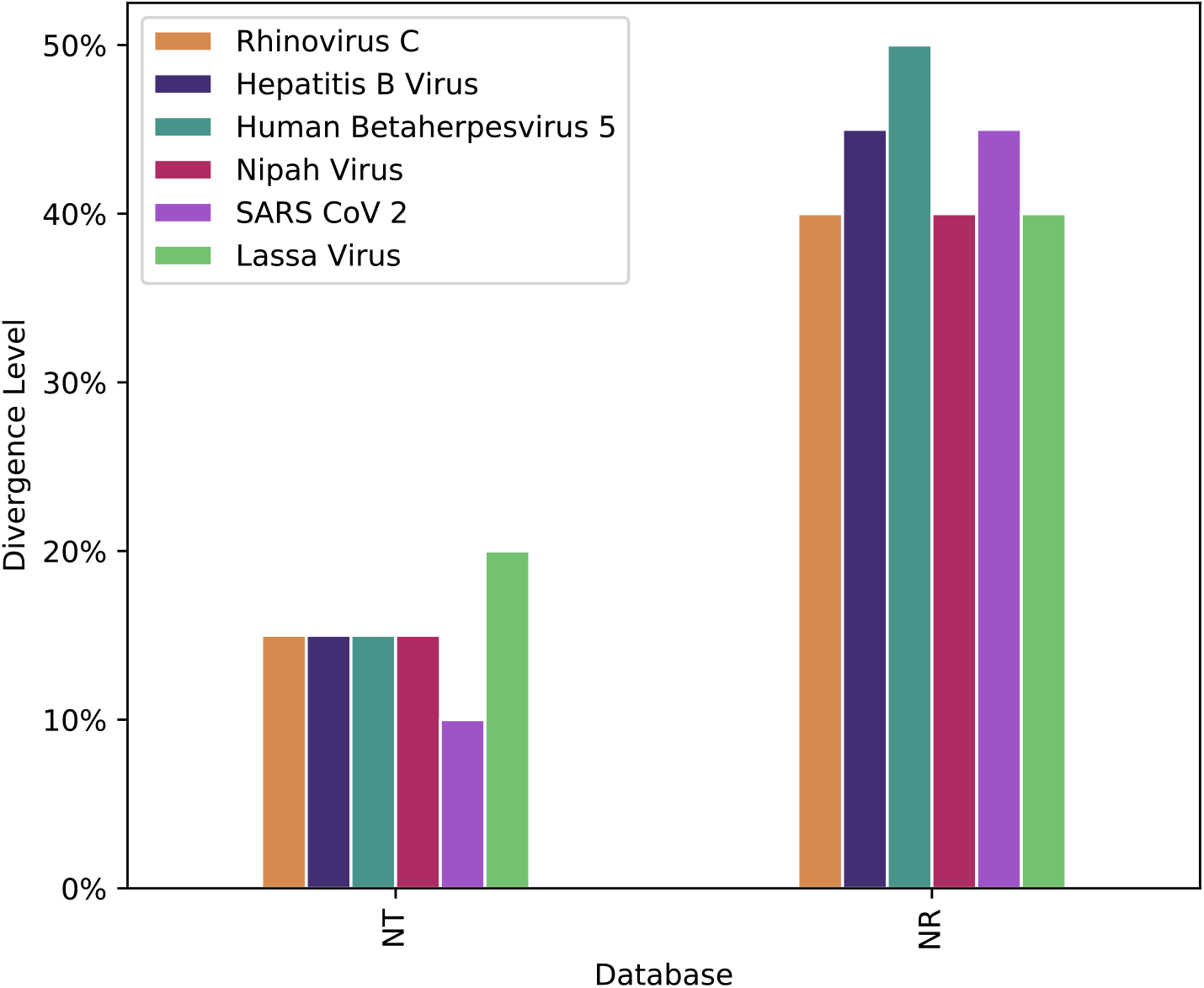
The maximum level of sequence divergence for viral detection on NT and NR databases for six viral species.

Overall, NR was able to detect sequences ≥ 20-30% more divergent than NT, suggesting an important role for protein alignments in expanding the detection of novel organisms (Fig. 5).

### Application II: Identification of microbes in human clinical samples

To determine the level of detection limit for a known RNA virus, HCoV OC43 virus, we generated samples containing varying levels of the multiplicity of infection (MOI), the ratio of virus particles to host cells (0.0001-1; Table 3) and ran them on CZ ID. The results showed that CZ ID could accurately detect HCoV OC43 virus at all MOI levels tested down to 0.0001 MOI, and the virus was not detected in the negative control (0 MOI). This demonstrated that CZ ID can detect known RNA viruses at varying abundance levels as a proxy for a human clinical sample, even at low abundances.

**Table 3.**
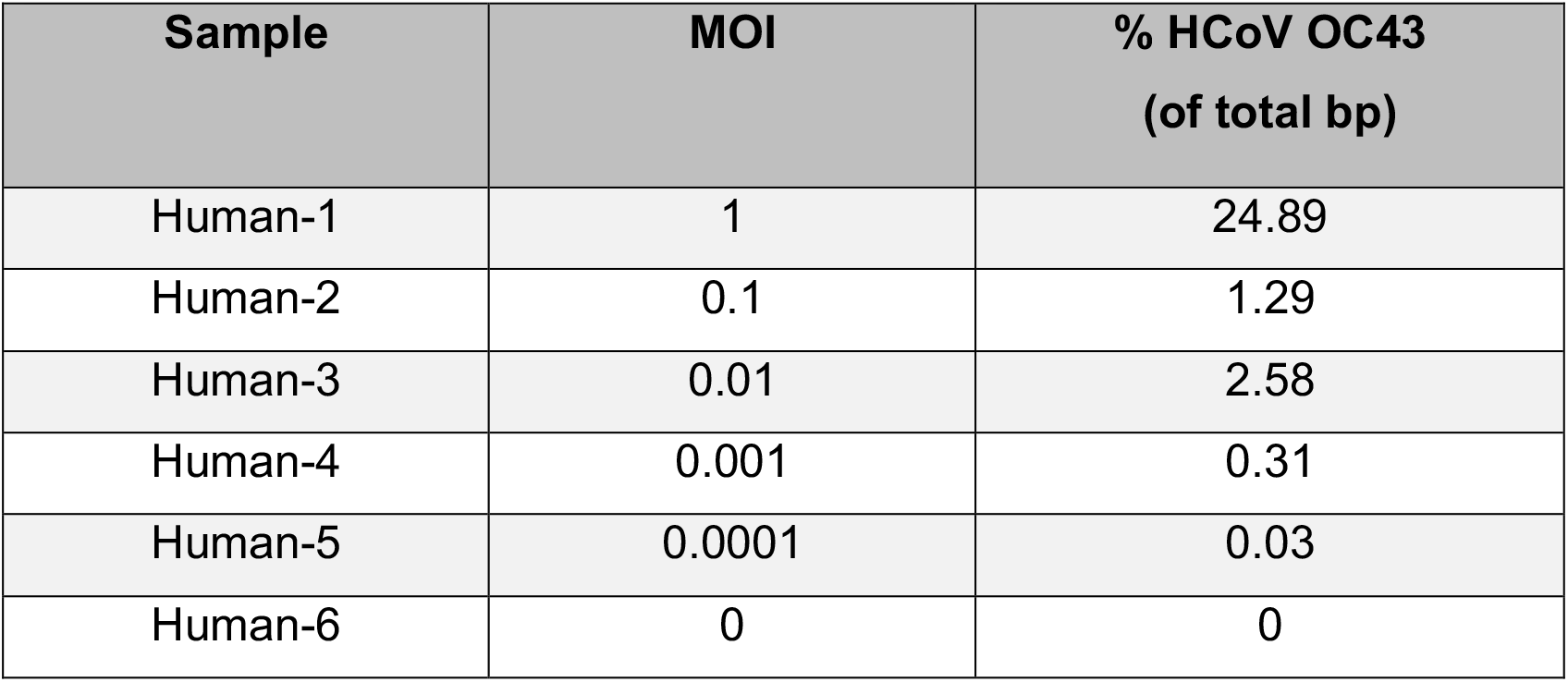
The abundance of virus detected using CZ ID from samples of HeLa cells infected with human coronavirus (HCoV OC43) at varying MOI, the ratio of virus particles to host cells. CZ ID detected HCoV OC43 virus down to 0.0001 MOI and not in the negative control.

| Sample | MOI | % HCoV OC43<br>(of total bp) |
| --- | --- | --- |
| Human-1 | 1 | 24.89 |
| Human-2 | 0.1 | 1.29 |
| Human-3 | 0.01 | 2.58 |
| Human-4 | 0.001 | 0.31 |
| Human-5 | 0.0001 | 0.03 |
| Human-6 | 0 | 0 |

### Application III: Identification of microbes in non-human host samples

We analyzed five orthogonally-characterized single mosquito samples to evaluate whether the pipeline could accurately identify known and novel viruses in complex metagenomic samples derived from non-human hosts (Batson *et al*., 2021).

After analyzing these samples with CZ ID, 66 hits to viruses were identified across the five samples (Table 4). We sought to orthogonally characterize each contig by using the BLASTn algorithm to search the most up-to-date NCBI standard nucleotide (NT/NR) database (query date 12/2023). This analysis confirmed 48 true positive hits: 21 where the top BLAST hit matched the initial CZ ID identification, and 27 where the top BLAST hit differed due to differences in the underlying databases but was consistent with the viruses identified in the previously published orthogonal samples (Batson *et al*., 2021).

**Table 4.**
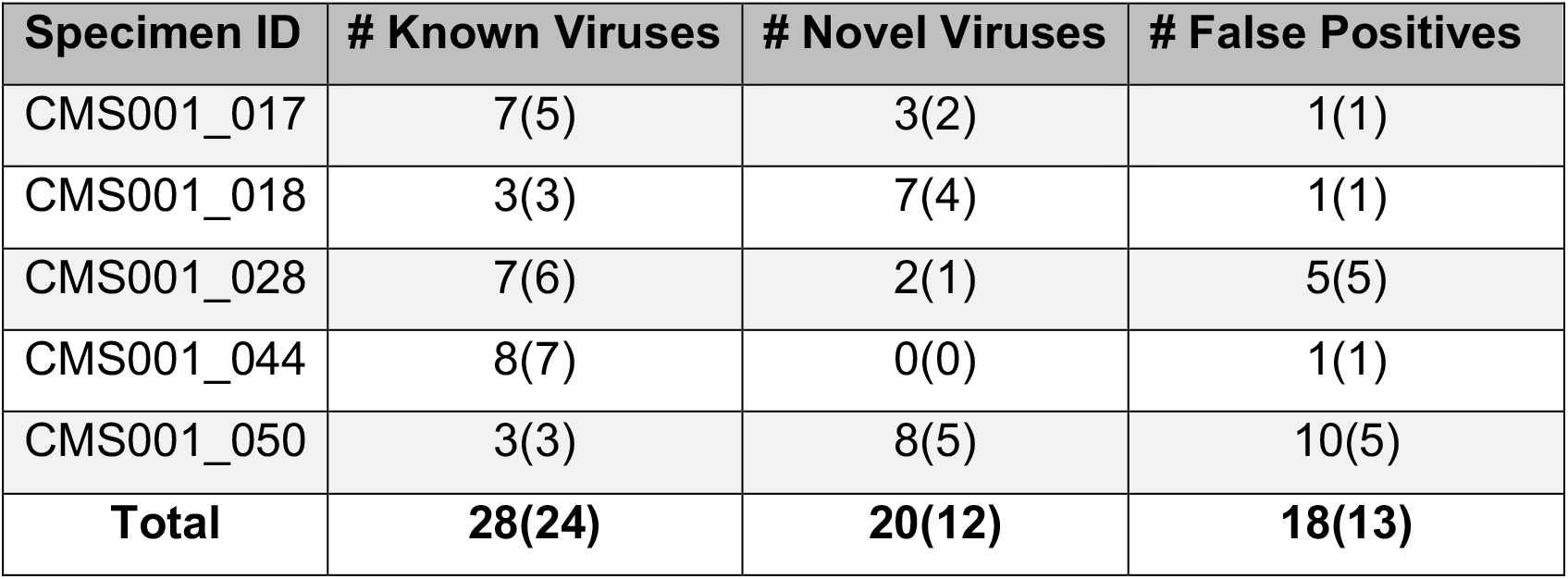
Total number of CZ ID viral hits (true positives) in known and novel viruses (with unique viral species confirmed by BLAST in parentheses) and false positives in samples derived from mosquitos.

| Specimen ID | # Known Viruses | # Novel Viruses | # False Positives |
| --- | --- | --- | --- |
| CMS001_017 | 7(5) | 3(2) | 1(1) |
| CMS001_018 | 3(3) | 7(4) | 1(1) |
| CMS001_028 | 7(6) | 2(1) | 5(5) |
| CMS001_044 | 8(7) | 0(0) | 1(1) |
| CMS001_050 | 3(3) | 8(5) | 10(5) |
| **Total** | **28(24)** | **20(12)** | **18(13)** |

A critical aspect of our investigation was the determination of false positives. Of the initial hits, we identified 18 as false positives, where downstream BLASTn analysis showed that the top hit was not a virus. Notably, most of these false positives (11 out of 18) aligned to sequences of mosquito species. Additionally, all viral hits below 210 bp alignment length to NR were false positives. Thus, setting an alignment length threshold greater than 210 bp effectively reduces the incidence of false positives in the analysis.

Excluding false positives, a total of 40 hits (83% of true positives) matched to viral species identified by Batson *et al*., (2021), highlighting the concordance of results obtained with CZ ID mNGS Nanopore pipeline and the results based on previously-published Illumina sequencing.

Our analysis also distinguished between novel and known virus species, identifying eight novel (20 total hits) and 15 known virus species (28 total hits). The downstream analysis using BLASTn showed that all viral hits considered to be known had NT % identity ≥ 88% via CZ ID, and viruses identified as novel had NT % identity ≤ 87% and NR % identity ≤ 74% via CZ ID. These results demonstrate the potential of CZ ID to support the detection and identification of emerging infectious diseases in diverse host species.

## DISCUSSION

Metagenomics is a powerful tool for studying infectious diseases, enabling the unbiased, direct detection and identification of pathogens. Due to its portability and low start-up costs, Nanopore sequencing has seen increased global adoption for local genomic pathogen surveillance since the COVID-19 pandemic (e.g., Tegally *et al*., 2022). However, using Nanopore sequencing for mNGS is an emerging technology in part because efficient analysis of the data remains challenging. Therefore, we have developed an easy-to-use pipeline that further unlocks the potential for researchers across the globe to use this technology for applications in infectious disease research regardless of computational power. This is especially important since the burden of infectious disease disproportionately impacts lower-middle-income countries (LMICs) (Marais *et al*., 2023).

We have described the pipeline implementation and discussed web app features to support data analysis. CZ ID performs equally well at estimating the relative abundance of a standard mock microbial community when benchmarked against a commonly used tool, Kraken2. CZ ID also detects divergent viruses using a simulated dataset, detects known viruses in clinical samples, and discovers known and novel RNA viruses in non-human hosts.

Individual bioinformatic tools for metagenomic analyses can be challenging to run, computationally expensive, and time-intensive. These issues scale when creating and maintaining pipelines consisting of multiple tools for comprehensive microbial analysis (from QC to assembly to alignment). The CZ ID platform automates validated bioinformatic pipelines so researchers can focus on using analysis results to inform the next steps of their projects. CZ ID is engineered to be fast and scalable, running on-demand and efficiently analyzing large numbers of samples concurrently and quickly, even against large databases (all of NCBI NT). The use of engineering best practices for dependency and error management ensures that results are reliable and consistent. The CZ ID support team provides continuing maintenance, updates, and user support.

One limitation of our study is its narrow performance evaluation compared to other tools. CZ ID is one of only a few tools that combine multiple aspects of analysis into a robust mNGS pipeline for Nanopore sequencing data (e.g., Fan *et al*., 2021), limiting the set of truly comparable tools. We focused our efforts on demonstrating a couple of relevant evaluations, including the ability to identify the relative abundance of known organisms and sensitivity for novel virus detection. The challenges associated with benchmarking (including the impact of tool selection, parameterization, databases, and datasets on the final metrics) are well-recognized. We hope that CZ ID’s ease of use encourages researchers interested in applying Nanopore mNGS to their research questions to perform additional and more comprehensive use-case-relevant benchmarking studies. One limitation of the current CZ ID implementation is its reliance on cloud infrastructure to provide ease-of-use benefits. Some scientific use cases require running software locally. For researchers with computational expertise and resources, the open availability of the CZ ID workflows enables the opportunity to run it offline.

CZ ID enables researchers with limited time or computational expertise to leverage insights from Nanopore data without setting up and maintaining their own pipeline. This makes the CZ ID mNGS Nanopore pipeline particularly well-suited for research and training in resource-limited settings, such as LMICs. Moreover, given the flexibility of CZ ID to accept sequencing data from any sample type or host organism, we look forward to seeing how CZ ID is applied to a broad array of research questions and benchmarked against a range of tools. We have already seen the CZ ID mNGS Nanopore pipeline applied to detecting microbes in rare animal species using eDNA (Koda *et al*., 2023).

## Acknowledgments

Thank you to Dr. Amy Kistler for providing the mosquito samples for sequencing and analysis. Figure 1 was created with BioRender.com.

## Supporting source code

All the CZ ID code is open-source and available at https://github.com/chanzuckerberg/czid-workflows/tree/main.

## Data availability

CZ ID results are available as a public project (Simmonds_et_al_2024_ONT_v1) at czid.org. The data sets generated to support the results of this article are available in the Sequence Read Archive (SRA) repository, [will cite unique persistent identifier once submitted to SRA].

## CZ ID Team author list

### Engineering

Julie Han^1^, Neha Chourasia^1^, Nina Bernick^1^, Omar Valenzuela^1^, Robert Aboukhalil^1^, Vincent Selhorst-Jones^1^*Product* Ann E. Jones^1^, Elizabeth Fahsbender^1^, Janeece Pourroy^1^, Kevin Wang^1^, Olivia Homes^1^, Samantha Scovaner^1^, Xochitl Butcher^1^

