## Supplementary material for "CZ ID: a cloud-based, no-code platform enabling advanced long read metagenomic analysis": Supp. Table 1

**Suppl. Table 1.** Details of the virus families, species, genome length and corresponding NCBI accession ID of the sequences used for simulating divergent viruses.

| Family | Species | Genome length (bp) | NCBI Accession IDs |
|---|---|---|---|
| Orthoherpesviridae | Human betaherpesvirus 5 (strain Merlin) (DNA) | 235,646 | NC_006273.2 |
| Hepadnaviridae | Hepatitis B virus (strain ayw) (DNA) | 3,182 | NC_003977.2 |
| Arenaviridae | Lassa virus, L segment (ss-RNA)<br>Lassa virus, S segment (ss-RNA) | 7,279<br>3,402 | NC_004297.1<br>NC_004296.1 |
| Paramyxoviridae | *Henipavirus nipahense* (ss-RNA) | 18,246 | NC_002728.1 |
| Picornaviridae | Human rhinovirus C (RNA) | 7,099 | NC_009996.1 |
| Coronaviridae | Severe acute respiratory syndrome coronavirus 2 isolate Wuhan-Hu-1 (ss-RNA) | 29,903 | NC_045512.2 |
