## Supplementary material for "CZ ID: a cloud-based, no-code platform enabling advanced long read metagenomic analysis": Supp. Table 2

**Supp. Table 2.** Details of the mosquito specimens sequenced in this study. Specimen were originally collected and analyzed with other methods in Batson *et al*., (2021).

| Specimen ID | Genus | Species | Sex | Collection date | Latitude | Longitude | Blood fed status | RNA concentration, ng/uL |
|---|---|---|---|---|---|---|---|---|
| CMS001_017 | *Culex* | *erythrothorax* | female | 10/6/17 | 37.5819 | -122.0484 | blood fed | 73.9 |
| CMS001_018 | *Culex* | *erythrothorax* | female | 9/27/17 | 37.557 | -122.0794 | blood fed | 46.4 |
| CMS001_028 | *Culex* | *tarsalis* | female | 9/12/17 | 37.7152 | -122.1943 | blood fed | 96 |
| CMS001_044 | *Culex* | *pipiens* | female | 8/25/17 | 37.6613 | -122.1334 | blood fed | 62 |
| CMS001_050 | *Culex* | *erythrothorax* | female | 10/17/17 | 37.5061 | -121.9986 | blood fed | 57 |
